# *Klf4*-dependent pivotal role of smooth muscle-derived *Gli1*^+^ lineage progenitor cells in lung cancer progression

**DOI:** 10.1101/2025.05.30.657094

**Authors:** Sizhao Lu, Andre C. Navarro, Amber M. Johnson, Tysen Noble, Austin J Jolly, Allison M Dubner, Karen S Moulton, Raphael A Nemenoff, Mary CM Weiser-Evans

**Affiliations:** Department of Medicine, Division of Renal Diseases and Hypertension, University of Colorado Anschutz Medical Campus, Aurora, CO, USA; Medical Scientist Training Program, University of Colorado School of Medicine, Anschutz Medical Campus, Aurora, CO, USA; Biomedical Sciences and Biotechnology MS program, University of Colorado Graduate School, Anschutz Medical Campus, Aurora, CO, USA; Department of Medicine, Division of Cardiology, University of Colorado Anschutz Medical Campus, Aurora, CO, USA; School of Medicine, Consortium for Fibrosis Research and Translation, University of Colorado Anschutz Medical Campus, Aurora, CO, USA; Cardiovascular Pulmonary Research Program, University of Colorado Anschutz Medical Campus, Aurora, CO, USA

## Abstract

Lung cancer remains the leading cause of cancer-related deaths worldwide. Despite the remarkable efficacy of immune checkpoint inhibitors, only a subset of lung adenocarcinoma (LUAD) patients respond and many eventually progress, underscoring the need to develop novel combination therapies. The interaction between cancer cells and the tumor microenvironment (TME), a complex niche including cells of the innate and adaptive immune system, cancer-associated fibroblasts (CAFs), vascular cells, and extracellular matrix (ECM) plays a vital role in tumorigenesis and cancer progression. Using fate mapping, we previously identified a novel subpopulation of resident vascular stem cells derived from vascular smooth muscle cells, designated AdvSca1-SM (vascular Adventitial location, Stem cell antigen-1 expression, SMooth muscle origin) cells. AdvSca1-SM cells are the predominant cell type responding to vessel wall dysfunction and their selective differentiation to myofibroblasts promotes vascular fibrosis. SMC-to-AdvSca1-SM cell reprogramming is dependent on SMC induction of the transcription factor KLF4, and KLF4 is essential for maintenance of the stem cell phenotype. The function of AdvSca1-SM cells in LUAD tumorigenesis has not been explored. Using an orthotopic immunocompetent mouse model of LUAD, we demonstrate that AdvSca1-SM cells are a major component of lung tumors, significantly contributing to cancer associated fibroblasts (CAFs). Compared to AdvSca1-SM cells, CAFs have altered ECM gene expression with a reduction of a stemness signature. AdvSca1-SM-specific genetic ablation of *Klf4* altered their phenotype resulting in inhibition of communication between cancer cells and CAFs, decreases in innate immunosuppressive populations, and increased T cell infiltration into the tumor. Targeting this population may represent a novel strategy to improve the response to immunotherapy.

## INTRODUCTION

Lung cancer remains the leading cause of cancer-related deaths worldwide in both men and women^1^. While the development of immunotherapy approaches has shown remarkable efficacy, only a subset of lung adenocarcinoma (LUAD) patients respond to this approach, and many eventually progress, underscoring the unmet need to develop novel combination therapies^2–5^.

Developing new therapeutic strategies requires a better understanding of the complex interactions between cancer cells and the tumor microenvironment (TME), a complex niche including cells of the innate and adaptive immune system, cancer-associated fibroblasts (CAFs), vascular cells, and extracellular matrix (ECM)^6–8^. While the role of cancer stem cells to cancer progression has been actively studied, the contribution of stem/progenitor populations in the stroma has been an understudied area, and represents an essential, but underappreciated contribution to the TME.

The blood vessel wall is a complex, multilayered tissue consisting of multiple cell types that dynamically communicate with each other to regulate vessel homeostasis and disease progression. The middle media layer consists of vascular smooth muscle cells (SMCs) that express high levels of SMC-specific contractile proteins necessary for their function in the maintenance of blood vessel homeostasis^9^. In disease, however, SMCs are capable of undergoing profound phenotypic and functional changes. The outer adventitial vessel layer is populated by dynamic populations of leukocytes, microvessels, fibroblasts, adipocytes, and resident progenitor cells^10–12^. These cell populations not only maintain normal vessel homeostasis, but also respond robustly to many kinds of vascular injury. The discovery that resident Stem Cell Antigen-1 (Sca1)(+) vascular progenitor cells (AdvSca1 cells) with multiple fate potentials reside in the adventitia raised new and important questions about these cells in growth, remodeling, repair, and disease^13–17^. Our previous report using a highly specific SMC lineage-mapping approach^18^ demonstrated that mature SMCs migrate into the adventitia and are reprogrammed into a distinct subset of AdvSca1 progenitor cells that we termed **AdvSca1-SM cells** (<u>Adv</u>entitial location, <u>SCA1</u>-positive, <u>S</u>mooth <u>M</u>uscle origin)^19^. AdvSca1-SM cells reside in multiple vascular beds (e.g. aorta, carotid and femoral arteries, and the pulmonary and coronary arterial vasculature)^19^. Induction of the pluripotency-associated transcription factor KLF4 is essential for SMC reprogramming to AdvSca1-SM cells and to the maintenance of the AdvSca1-SM stem cell phenotype^19^. AdvSca1-SM cells exhibit a multipotent phenotype marked by their ability to differentiate into multiple cell types in response to specific environmental cues^19^. Importantly, these cells are uniquely poised to rapidly respond to injury and contribute to disease progression; our recent studies demonstrated that AdvSca1-SM cells are activated in response to acute and chronic vascular disease, differentiate preferentially to myofibroblasts, and are major contributors to vascular injury disease-associated adventitial remodeling, fibrosis, and plaque formation. The contribution of AdvSca1-SM cells to the lung cancer TME and their role in lung cancer progression and metastasis are entirely unknown.

Here, using a highly selective AdvSca1-SM lineage approach and a syngeneic orthotopic mouse model of LUAD, we show that AdvSca1-SM cells localize in pulmonary artery and airway adventitial niches in normal tumor-free lungs. In response to tumor formation and progression, AdvSca1-SM cells expand in number, preferentially differentiate into CAFs, and contribute to tumor formation and metastasis. Surprisingly, AdvSca1-SM-specific deletion of *Klf4* resulted in smaller primary tumors with no secondary pulmonary metastases. This was associated with an alteration of the AdvSca1-SM cell transcriptomic profile as well as increased T cell tumor infiltration and promotion of an anti-tumorigenic macrophage/monocyte phenotype. Our findings indicate that lung AdvSca1-SM cells are major regulators of tumor progression and that a shift in AdvSca1-SM cell phenotype through *Klf4* deletion blunts tumor progression through alterations in the TME, thus highlighting this population as a potential therapeutic target.

## MATERIALS AND METHODS

### Mice

*Gli1*-Cre^ERT^ (JAX stock 007913) and ROSA26-YFP reporter mice (JAX stock 006148) were obtained from Jackson Laboratory. *Klf4* floxed mice were obtained from Dr. Klaus H. Kaestner (University of Pennsylvania). All mice were fully backcrossed to a C57BL/6 genetic background prior to studies. *Gli1*-Cre^ERT^ transgenic mice and Rosa26-YFP reporter mice were bred to generate tamoxifen-inducible AdvSca1-SM cell-specific YFP-expressing reporter mice (*Gli1*-Cre^ERT^-YFP). *Gli1*-Cre^ERT^-YFP mice were bred to *Klf4* floxed mice to generate AdvSca1-SM cell-specific *Klf4* knockout mice (*Gli1*-Cre^ERT^-YFP;*Klf4*^flox/flox^). Male and female 8-week old *Gli1*-Cre^ERT^-YFP and Gli1-CreERT-YFP;Klf4flox/flox mice received 1 mg IP tamoxifen injections for 12 consecutive days to induce YFP reporter knock-in and *Klf4* knockout prior to experiments. The tamoxifen was allowed to wash out for 5 days prior to tumor cell injection and initiation of experiments.

### Orthotopic lung cancer model

Primary tumors were generated by implanting murine lung cancer cells directly into the lungs of syngeneic *Gli1*-Cre^ERT^-YFP and *Gli1*-Cre^ERT^-YFP;*Klf4*^flox/flox^ mice as previously described^20,21^. Briefly, cells were suspended in Hank’s Balanced Salt Solution (HBSS; Corning) containing 1.35 mg/mL Matrigel Basement Membrane Matrix (Corning; #354234). Mice were anesthetized, a skin incision was made along the left lateral axillary line, subcutaneous fat was removed to visualize the left lung, and the cell suspension was injected directly into the left lung parenchyma using a 30G needle. The incision was closed using veterinary skin adhesive or staples. Cells were injected at the concentration of 1x10^5^ (LLC; KRas^G12C^ mutation) or 2x10^5^ (CMT167; KRas^G12V^ mutation) cells per mouse. Mice were harvested 3 (LLC) or 4 (CMT167) weeks after injections.

### Immunofluorescent staining and analysis

Harvested tumor tissues were fixed in 10% buffered formalin phosphate (Fisher Scientific) for 48 hours and stored in 70% ethanol for 8-24 hours until processing and paraffin embedding. Tissues were sectioned at 5µm thickness with a microtome. Sections were deparaffinized through a series of xylene and graded ethanol steps, underwent antigen retrieval (Vector) followed by blocking in normal horse serum (Vector) Sections were then incubated overnight at 4°C with a rabbit monoclonal anti-GFP antibody (1:200, Abcam, Cat #Ab290) and/or a rat anti- CD3 antibody (1:100; Bio-Rad, Cat #MCA1957) followed by corresponding Alexa Fluor 488- and Alexa Fluor 654-conjugated secondary antibodies (1:500; Invitrogen) and an anti-αSMA-Cy3 antibody (Sigma Aldrich, Cat #C6198, 1:2000) for 60 minutes at room temperature. Sections were mounted with Vectashield mounting media with DAPI (Vector Laboratories, Cat #H-1200). Images were captured with a Keyence BZ-X710 fluorescent microscope. Cellpose3 was used for fluorescent image segmentation. The pretrained model CP was further trained with a set of randomly selected 4 images. All images were segmented using the custom model and identified cells were exported as imageJ ROIs with the parameter --save_rois. ImageJ macro was used to quantify green and red fluorescence in all the ROIs. Custom python code was used to identify YFP+ and CD3+ cells. The identified positive cells were exported as ImageJ ROI and visually examined to rule out any false positive due to autofluorescence. Final quantification results were averaged per sample and expressed as cell count per 40x field.

### scRNA-seq

Single cell suspension was made of the tumor bearing and naïve non-tumor bearing left lungs of *Gli1*-Cre^ERT^-YFP and *Gli1*-Cre^ERT^-YFP;*Klf4*^flox/flox^ mice. Harvested tissues were minced and digested in 10 mL warm digestion buffer (3.2 mg/mL collagenase II, 0.7 mg/mL elastase, 0.2 mg/mL soybean trypsin inhibitor) at 37°C for 1 hour. Lung tissues were physically disrupted using a 10 mL serological pipet every 10 minutes during the digest. Cell suspension was filtered through a 250 μm filter, washed with FA3 buffer (1x PBS, 1mM EDTA, 25mM HEPES (pH 7.0), 1% FBS) and filtered through a 70 μm filter. Cell suspensions were pelleted and resuspended in 500 uL of FA3 buffer. DAPI was added to the final concentration of 10 ng/mL for live/dead staining. Fluorescence activated cell sorting (FACS) was performed to separate YFP^+^ and YFP^-^ live cells. Sorted samples were subjected to scRNA-seq using the Chromium Single Cell 3’ Library and Gel Bead Kit (v3.1 10x Genomics, Pleasanton, CA) and the Chromium X. Libraries were sequenced on an Illumina Novaseq 6000 at Genomics Shared Resource at the University of Colorado Anschutz Medical Campus. 50,000 reads per cell were obtained. Fastq files were aligned to custom build GRCm39 reference (Ensembl, r104) with the addition of YFP gene, using Cell Ranger 6.1.2. Scanpy 1.9.3^22^ was used for the downstream analysis including quality control, normalization, clustering, based on Single-cell best practices^23^. Cells with less than 200 genes and genes expressed by less than 10 cells were filtered out. Cells with a percentage of mitochondrial counts exceeding 8% or with more than 7500 genes were filtered. Pseudobulk profiles were created from the single-cell data using decoupler 1.6.0^24^. Differential gene expression analysis was performed with python implementation of the framework DESeq2^25^.

Functional enrichment of biological terms was performed with decoupler 1.6.0 using GO_Biological_Process_2021.gmt, Reactome_2022.gmt, and KEGG_2019_Mouse.gmt obtained from Enrichr gene-set library^26^. Transcription factor activity inference was performed with decoupler 1.6.0 using CollecTRI gene regulatory network data^27^. LIANA+^28^ was used to infer steady-state ligand-receptor interactions and Tensor-Cell2cell^29^ was used for Intercellular Context Factorization based on official tutorials.

### Data availability

The NCBI Gene Expression Omnibus database accession number for the data reported in this paper is GSE269657.

### Statistical Analysis

All experiments reported were carried out with at least 3 biological replicates, including both male and female mice. Data from male and female mice were combined for analysis. Data were analyzed using GraphPad Prism 10 (GraphPad Software, Inc). Shapiro-Wilk test (n≥3) or D’Agostino & Pearson test (n≥8) were performed to determine the normality of the data. Brown-Forsythe test was used to examine the equality of group variances. One-way ANOVA with Bonferroni’s post-hoc test was used to compare between multiple groups. For data that failed the normality test or equal variances test, a Kruskal-Wallis test was used to compare the groups followed by Dunn’s multiple comparison tests. Histochemical- and immunofluorescently stained sections were visually examined and only sections with high quality tissue and staining were included in the quantitative analysis. P-values <0.05 were considered statistically significant.

## RESULTS

An immunocompetent orthotopic mouse model of lung cancer progression and metastasis developed in our lab was used to examine the contribution of AdvSca1-SM cells to tumor formation. Prior to cancer cell implantation, mice received tamoxifen injections to induce YFP expression selectively in AdvSca1-SM cells (**Figure 1A**). Please note that tamoxifen treatment prior to the start of the experiment results in efficient, permanent, and selective YFP knockin only in AdvSca1-SM cells^30–32^. As tamoxifen is washed out prior to tumor cell implantation, the only cells expressing YFP at the time of tissue harvest are AdvSca1-SM cell-derived. Even if *Gli1* is induced in other cell types during tumor formation, they will not be labeled with YFP as tamoxifen is not present in the system to activate Cre recombinase and thus knock in YFP. After tamoxifen washout LLC cancer cells harboring a KRas^G12C^ mutation were directly injected into the left lung of syngeneic AdvSca1-SM lineage mice, therefore subjecting cancer cells to the correct microenvironment. Naïve control and tumor-bearing lung tissues were harvested at 3 weeks post-implantation. Immunofluorescent (IF) staining for the YFP AdvSca1-SM lineage mark and αSMA was performed to visualize YFP^+^ AdvSca1-SM and AdvSca1-SM-derived cells in tissue sections. Consistent with our previous reports^19,30–32^, *Gli1* lineage AdvSca1-SM cells exhibited a perivascular distribution in control non-tumor-bearing lung tissue (**Figure 1B**, top panel). Interestingly, YFP^+^ AdvSca1-SM cells were also detected around airways in close association to the airway αSMA^+^ smooth muscle layer, suggesting the intriguing possibility of reprogramming of airway smooth muscle cells to progenitor cells, similar to vascular smooth muscle cell-to-AdvSca1-SM cell reprogramming we reported previously^19^. Note that the only YFP+ cells at the start of the experiment were located in these perivascular and peri-airway niches. In tumor bearing lungs, we found large numbers of YFP+ AdvSca1-SM-derived cells within the core of the primary tumor (**Figure 1B**, bottom panel), supporting their role in facilitating tumor progression and metastasis. This finding was consistent in all tumor-bearing lungs examined (N=6, **not shown**). In addition, injection of a second cancer cell type, CMT167 harboring a KRas^G12V^ mutation, also induced expansion and infiltration of AdvSca1-SM cells within the core of the primary tumor and surrounding the periphery of the tumor (**Supplemental Figure I**).

**Figure 1.**
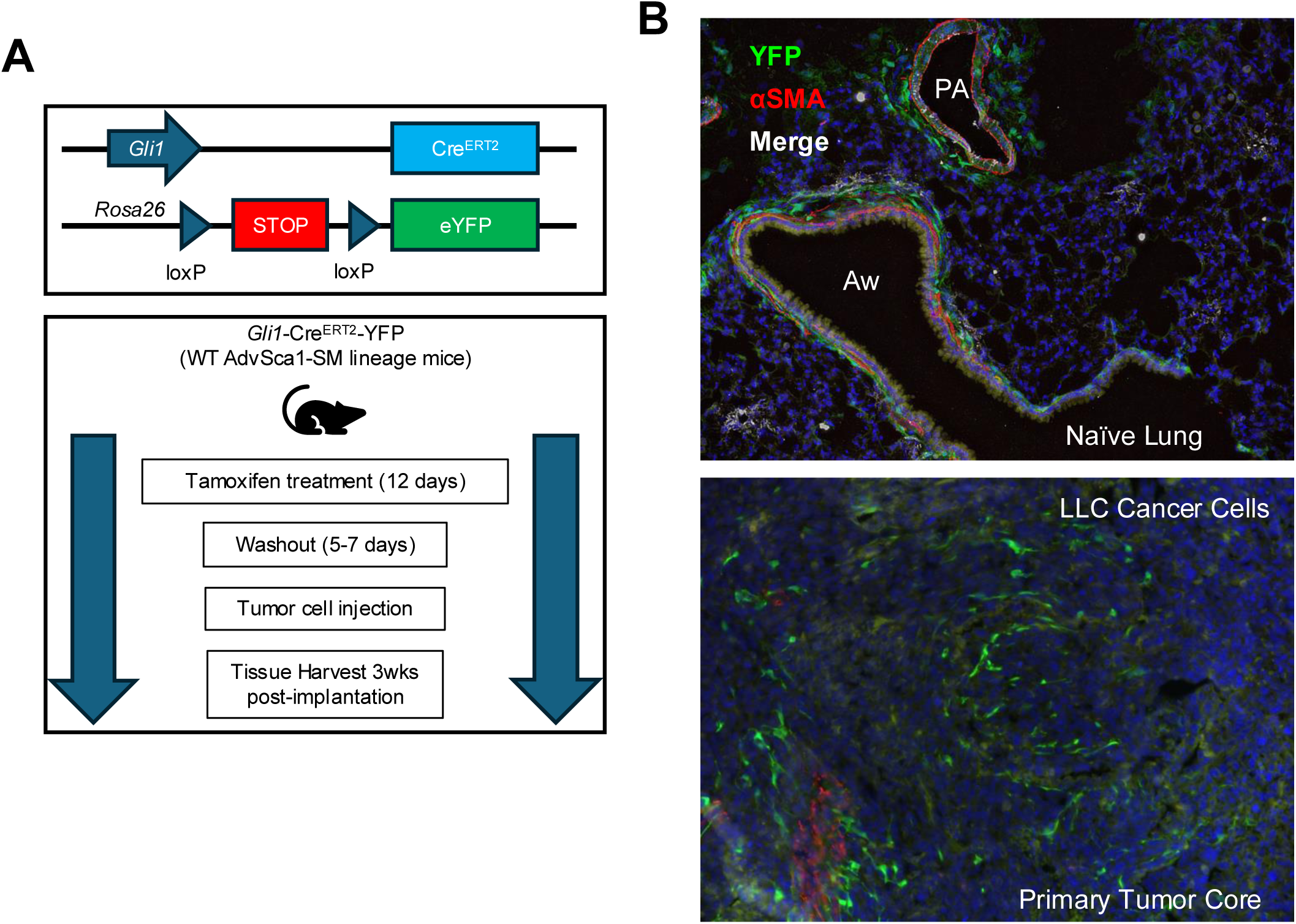
AdvSca1-SM cells reside in the lung and contribute to lung cancer progression and metastasis. **(A)** Schematic illustrating the experimental design. *Gli1*-Cre^ERT^-YFP mice were treated with tamoxifen to induce YFP gene knock-in specifically in AdvSca1-SM cells. After a washout period, mice were injected with LLC cancer cells orthotopically into the left lobe of the lung. **(B)** Lung tissues from naïve non-tumor-bearing **(top)** and LLC tumor-bearing mice **(bottom)** were immunofluorescently stained for AdvSca1-SM-derived cells (YFP, green), alpha smooth muscle actin (aSMA, red) for vascular and airway smooth muscle, and DAPI to identify cell nuclei. PA = pulmonary artery; Aw = airway. Representative 40x images shown from N=6.

To define the spectrum of *Gli1* AdvSca1-SM lineage cell phenotypes in the setting of tumorigenesis, we performed scRNA-seq with YFP-positive AdvSca1-SM-derived and YFP- negative non-AdvSca1-SM cells sorted from single cell suspension prepared from either naïve or LLC tumor-bearing lungs. Dimension reduction and clustering were performed to visualize the data (**Figure 2A**). Clusters were annotated using top expressing markers of each cluster (**Supplemental Figure IIA**), as well as mapping the data to the Human Lung Cell Atlas (HLCA, **Supplemental Figure IIB**)^33^. We selectively visualized the distribution of YFP-positive cells (**Figure 2B**) and their contribution to individual clusters (**Supplemental Figure IIC**). In naïve lung, the largest cluster (37.99%) of YFP-positive cells contributed to the annotated AdvSca1- SM cluster. Consistent with IF data (**Figure 1B**), fewer, but a significant portion of YFP-positive cells were present in the clusters of resident lung fibroblast populations (AlvFib_1: 20.92%, AlvFib_1: 10.83%, and BroncFib 20.77%). YFP-positive cells were also observed in smooth muscle cell (SMC) populations (7.66%), suggesting a dynamic interchange between SMCs and AdvSca1-SM cells in lung tissue. The presence of a lung tumor drastically altered the phenotype of lung YFP-positive cells (**Figure 2C & Supplemental Figure IIC**) with the great majority (86.75%) of YFP-positive cells adopting a cancer associated fibroblast (CAFs) phenotype.

**Figure 2.**
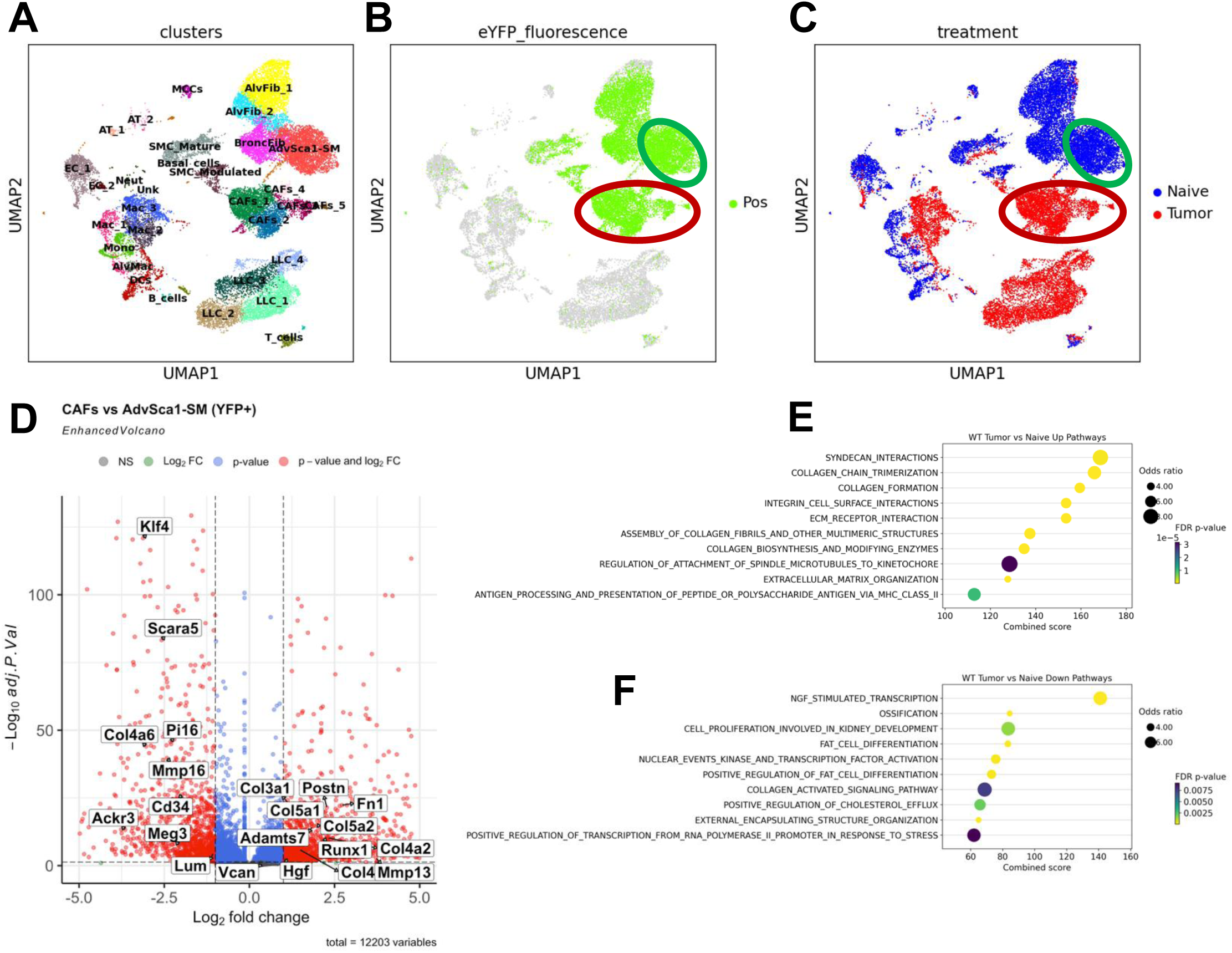
AdvSca1-SM lineage cells predominately contribute to cancer-associated fibroblasts (CAFs) in the tumor microenvironment (TME). Single cell RNA-sequencing (scRNA-Seq) was performed with single cell suspensions from naïve and LLC tumor-bearing lungs harvested from *Gli1*-Cre^ERT^-YFP AdvSca1-SM lineage mice. YFP^+^ AdvSca1-SM lineage cells and YFP^-^ cells were separated using FACS and subjected to scRNA-seq. UMAPs with annotated cell clusters **(A)**, highlighting YFP^+^ samples in green **(B),** and separating naïve and tumor samples **(C)** are shown. **(D)** Differential gene expression (DGE) analysis was performed between YFP^+^ CAFs (**red circle, panels B&C**) and AdvSca1-SM cells (**green circle, panels B&C**). The results are shown in the volcano plot with select genes labeled. Positive logFC indicate up-regulation in CAFs. Pathway enrichment for up-regulated genes **(E)** and down-regulated genes **(F)** in CAFs are shown.

From these data we conclude that the presence of cancer cells leads to expansion of AdvSca1-SM cells that contribute a major component of CAFs.

Differentiatial gene expression (DGE) analysis comparing YFP-positive CAFs to AdvSca1-SM cells showed that CAFs exhibited increased ECM related gene expression including expression of *Postn*, a well-known marker of activated myofibroblasts^34^ (**Figure 2D&E**). A wide range of additional ECM related genes were up-regulated in CAFs (**Supplemental Table 2&3**), including Types I, III, IV, VI, and XI collagen, fibronectin, integrins, and proteoglycans. Among the down-regulated genes in CAFs, we identified genes related to stemness of AdvSca1-SM cells, including *Klf4*, *Scara5*, *Pi16*, *Cd34*, *Ackr3,* and *Meg3* (**Figure 2D**). Interestingly, select ECM-related genes and pathways (**Figure 2D&F**) were down-regulated in CAFs, including multiple A Disintegrin and Metalloproteinase with Thrombospondin motifs (Adamts) genes and Matrix Metalloproteinase (Mmp) genes (**Supplemental Table 2&4**), suggesting altered ECM turnover and remodeling dynamics with tumor progression. Basement membrane components showed varied expressions, with up-regulation of *Col4a1, Col4a2* and down-regulation of *Col4a3, Col4a5, Col4a6* (**Figure 2D, Supplemental Table 2, 3&4**), likely linked to altered basement membrane permeability and cancer cell invasion.

AdvSca1-SM cells represented a mere 1.72% of YFP-negative samples in the naïve lung (**Supplemental Figure IID**), supporting the efficiency and specificity of the *Gli1* AdvSca1- SM lineage tracing model in labelling of AdvSca1-SM cells. 17.32% of YFP-negative cells were alveolar fibroblasts (AlvFib)(**Supplemental Figure IID**). We found that the macrophage cluster 1 (Mac_1 10.49%), Monocyte cluster (Mono, 8.58%), alveolar macrophage cluster (AlvMac, 3.29%), and dendritic cells (DCs, 10.23%) were the major populations of immune cells in the YFP-negative in naïve lung (**Supplemental Figure IID**). As expected, LLC cell clusters represented the vast majority (66.08%) of YFP-negative cells in LLC tumor-bearing lungs (**Supplemental Figure IID**). To better compare the composition changes between the tumor and naïve lung, we further plotted the composition of YFP-negative non-tumor portions of the samples (**Supplemental Figure IIE**). Immune cell populations accounted for 86.45% of non-tumor YFP^-^ cells in tumor-bearing lungs. Tumorigenesis shifted the macrophage phenotype from Mac_1 to Mac_2 (25.36%) and Mac_3 (42.65%), while shrinking AlvMac (0.25%) and Mono (4.21%) clusters. DGE analysis comparing the tumor associated macrophages (TAM) and naïve macrophages and monocytes (**Supplemental Figure IIIA, Supplemental Table 5**) and pathway analysis (**Supplemental Figure IIIB&C, Supplemental Table 6&7**) indicated the up-regulation of glycolysis related genes/pathways and interferon response genes, with down-regulation of cytokine production and cell adhesion molecules, consistent with previously reported transcriptomics profiles of TAMs by our group^20^. Transcription factor (TFs) activity inference results based on transcriptomic changes are shown in **Supplemental Figure IIID)**.

We previously demonstrated that the transcription factor KLF4 is required for the reprogramming of SMC to AdvSca1-SM cells^19^ and the maintenance of the stem-like phenotype of AdvSca1-SM cells^30^. Further, our recent studies indicate that genetic AdvSca1-SM cell-specific *Klf4* depletion altered the response of AdvSca1-SM cells in atherosclerosis^32^ and cardiac fibrosis (manuscript under review) promoting a cardiovascular protective effect in these settings. Here we investigated the effect of genetic AdvSca1-SM cell-specific *Klf4* deletion in *Gli1* AdvSca1-SM lineage cells on LLC tumorigenesis. *Gli1*-Cre^ERT2^-YFP (WT) and *Gli1*-Cre^ERT2^- YFP-*Klf4* KO (*Klf4* KO) mice were subjected to orthotopic LLC tumor implantation. Strikingly, tumor burden was significantly reduced in *Klf4* KO mice (**Figure 3A&B**). In addition, whereas all *Klf4* WT mice exhibited secondary pulmonary metastases to other lung lobes, none of the AdvSca1-SM *Klf4* KO mice exhibited secondary pulmonary metastases (**Supplemental Figure IVA**), supporting that manipulating the phenotype of AdvSca1-SM cells through loss of KLF4 expression is sufficient to inhibit both tumor growth and metastasis.

**Figure 3.**
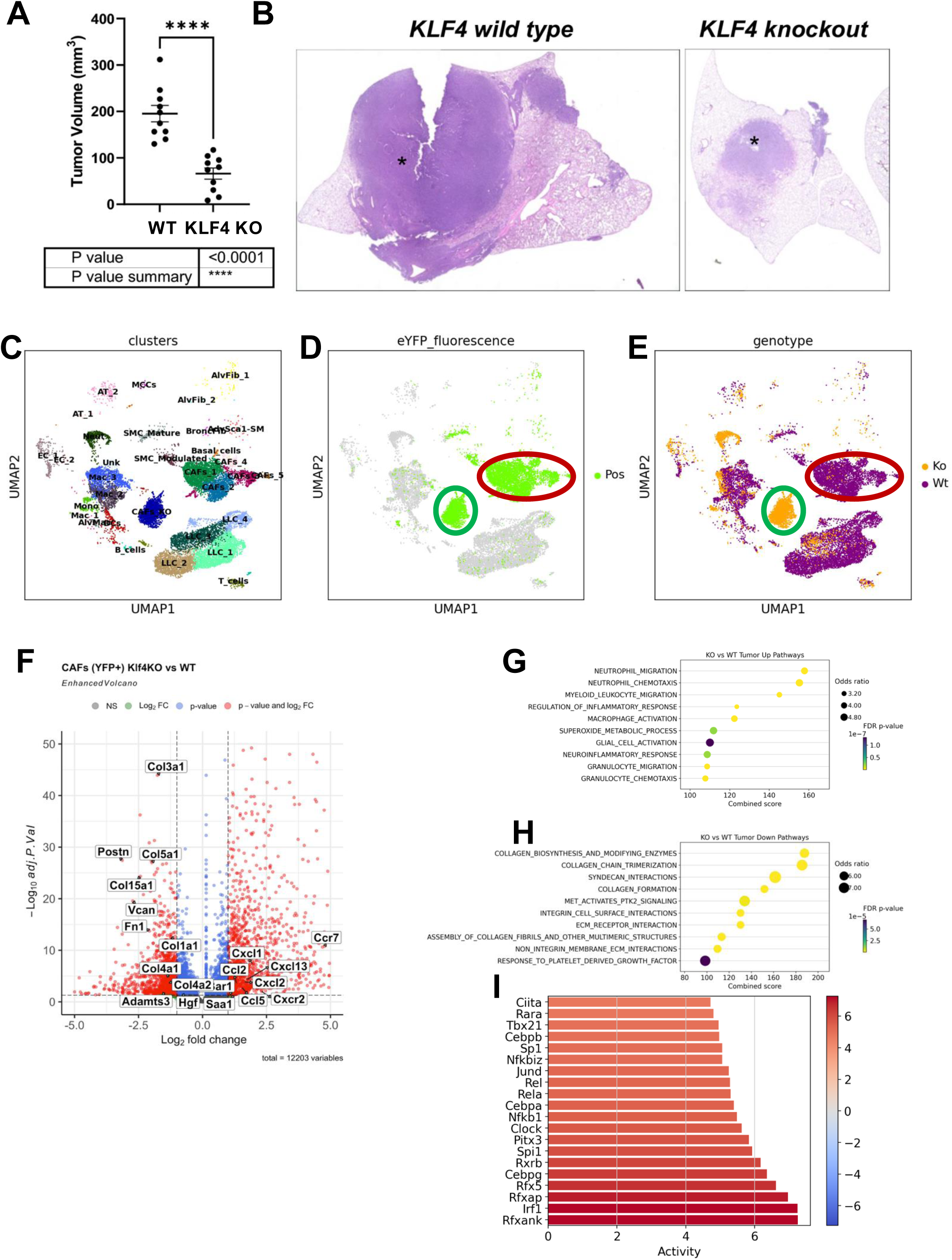
*Klf4* depletion in *Gli1*^+^ AdvSca1-SM lineage cells blunts tumorigenesis and alters CAF phenotype in the TME. Tamoxifen-treated *Gli1*-Cre^ERT^-YFP (WT) and *Gli1*-CreERT-YFP;*Klf4*^flox/flox^ (*Klf4* KO) mice were subjected to orthotopic LLC cell injection. Tumor volumes were measured by caliper **(A)** and H&E images (4x) of representative tumors from WT (**left**) and *Klf4* KO (**right**) mice are shown **(B)** scRNA-seq was performed with tumor bearing lungs from *Klf4* KO mice and the data integrated with WT scRNA-seq data. UMAP plots with annotated clusters **(C)**, highlighting YFP^+^ cells in green **(D)**, and separating WT and KO cells **(E)** are shown. **(F)** DGE analysis was performs comparing WT and *Klf4* KO YFP^+^ CAFs. Volcano plot shown with select genes labeled. Positive logFC indicate up-regulation in *Klf4* KO. Pathway enrichment results with up-regulated genes **(G)** and down-regulated genes **(H)** in *Klf4* KO CAFs are shown. **(I)** Transcription factors (TFs) activities were inferred based on the differential gene expression analysis. Barplot showing top activated (red) TFs.

Single cell suspensions from WT and *Klf4* KO LLC tumor-bearing lung tumors were subjected to scRNA-seq analysis to further examine the transcriptomic changes. In the annotated UMAP (**Figure 3C**), we identified a cluster of YFP-positive cells that are specific to the *Klf4* KO sample (CAFs_KO, **Figure 3D&E, green circle**), which is distinct from the WT CAF clusters (CAFs_1-5, **Figure 3D&E, red circle**). CAFs_KO cluster comprised 84.36% of the YFP-positive sample in *Klf4* KO lungs (**Supplemental Figure IVB**). DGE analysis comparing CAFs from *Klf4* KO samples with WT CAFs (**Figure 3F, Supplemental Table 8**) and pathway analysis (**Figure 3G&H, Supplemental Table 9&10**) showed that *Klf4* depletion caused the induction of cytokines that control immune cell migration and recruitment, and the suppression of ECM gene expression. TFs activity inference indicated the activation of proinflammatory TFs, such as NFkB, CiiTA, and IRF1. Further, RNA velocity analysis revealed the dynamic differentiation trajectory of AdvSca1-SM cells and CAFs (**Supplemental Figure IVC**). WT AdvSca1-SM-derived CAFs exhibited high RNA velocity length compared to naïve AdvSca1-SM cells (**Supplemental Figure IVD**), suggesting tumor progression induced rapid changes in gene transcription of WT AdvSca1-SM-derived CAFs. In addition, WT AdvSca1-SM-derived CAFs exhibited high velocity confidence compared to stemlike AdvSca1-SM cells from naïve lungs, suggesting they are moving through coherent differentiation trajectory (**Supplemental Figure IVE**). In contrast, *Klf4* KO CAFs exhibited low velocity length with high confidence, suggesting AdvSca1-SM-specific *Klf4* depletion blunted the transcription changes with AdvSca1-SM-derived CAFs likely entering a stable state (**Supplemental Figure IVC&D**).

We additionally examined the LLC clusters from WT and *Klf4* KO tumors. RNA velocity analysis showed trajectory originating from LLC_4 toward LLC_2 through LLC_3 and LLC_1 (**Figure 4A)**. Cell cycle analysis (**Figure 4B**) supported that LLC_4 has high G2M score (cell division) while part of LLC_1 had high G2M and S score (DNA replication). Interestingly, composition analysis showed that compared to WT, *Klf4* KO tumors exhibited reductions in LLC_1 and LLC_4 clusters, but a large increase in LLC_3 cluster (**Figure 4C**). In agreement, DGE analysis (**Figure 4D, Supplemental Table 11**) and pathway analysis (**Figure 4E&F, Supplemental Table 12&13**) showed that LLC cells from *Klf4* KO samples had reduced cell proliferation gene expression. The up-regulated genes are enriched in pathways related to immune responses and antigen presentation, suggesting a more immune-active tumor in *Klf4* KO LLC-injected lungs. TFs activity inference (**Figure 4G**) also supported activation of an immune response and a reduction in proliferation in *Klf4* KO samples, consistent with the reduced tumor size in *Klf4* KO mice.

**Figure 4.**
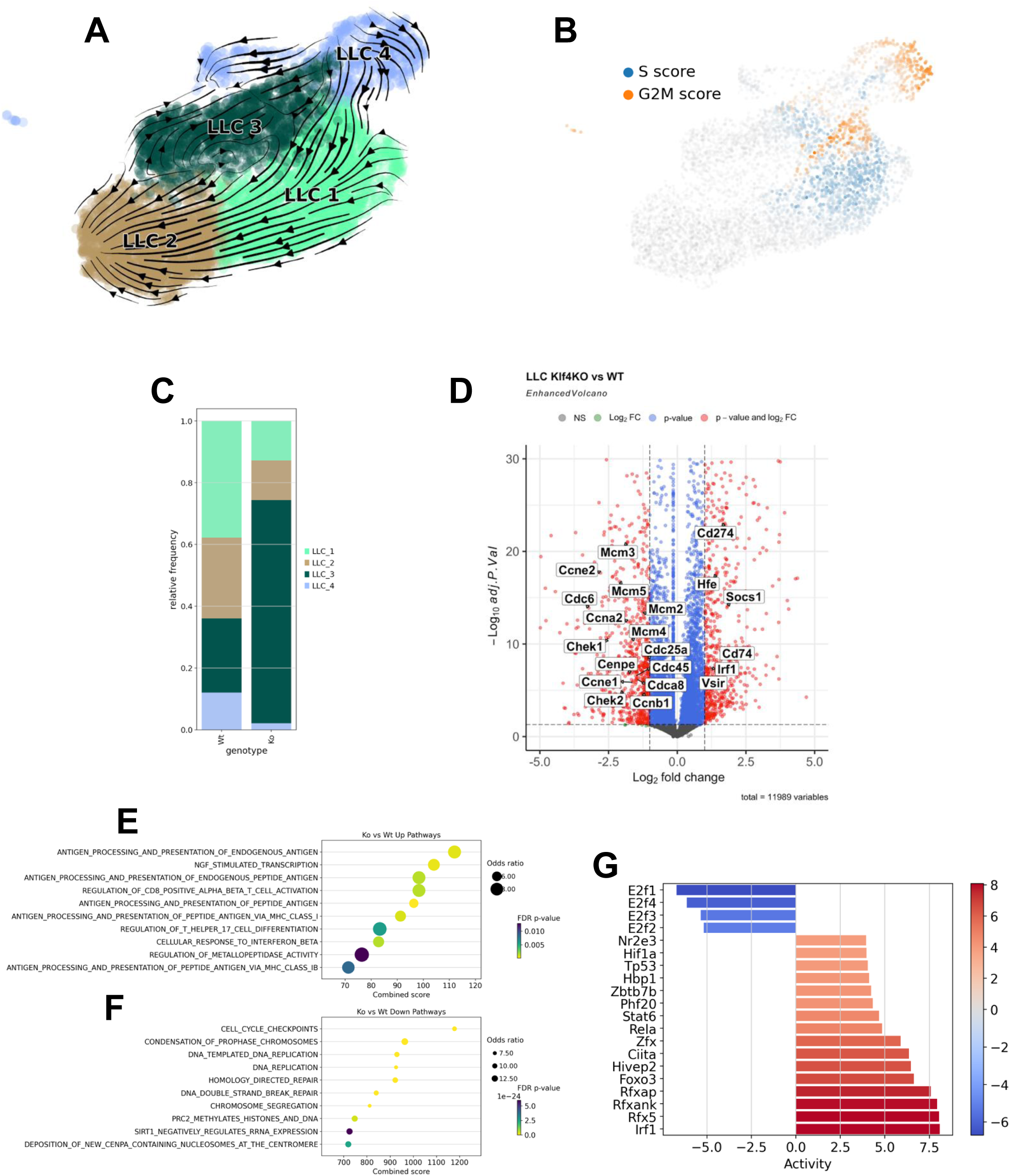
LLC tumor cells in *Klf4* KO lung are less proliferative and more immune active. RNA velocity analysis was performed for LLC cells in the scRNA-seq data. **(A)** RNA velocity is shown with stream plot. Arrows represent RNA velocity vectors, indicating the predicted future state of each cell. The length and direction of the arrows suggest the rate and trajectory of transcriptional changes, respectively. **(B)** LLC cells were scored for marker genes of S and G2M phase. UMAP colored by respective score are shown. **(C)** Composition analysis was performed to contrast LLC phenotype changes between WT and *Klf4* KO lungs. **(D)** DGE analysis was performed between LLC cells in WT and *Klf4* KO samples. Volcano plot with select genes labeled. Positive logFC indicate up-regulation in *Klf4* KO. Pathway enrichment with up-regulated genes **(E)** and down-regulated genes **(F)** in *Klf4* KO are shown. **(G)** Transcription factor (TFs) activities were inferred based on the differential gene expression analysis. Barplot shows activated (red) and suppressed (blue) TFs in LLCs from *Klf4* KO lungs.

Analysis of CAFs and LLC clusters supported that AdvSca1-SM-specific *Klf4* KO alters the TME to favor immune cell recruitment and anti-tumor responses. Therefore, we focused on the immune cell populations in the scRNA-seq data. Composition analysis of YFP-negative samples (**Supplemental Figure VA**) and YFP-negative cells excluding LLC tumors (**Supplemental Figure VB**) showed that, compared to WT, *Klf4* KO samples had reduced macrophage populations (WT: 68.39%, KO: 36.89%) and in particular reduced Mac_2 (WT: 25.36%, KO: 17.53%) and Mac_3 (WT: 42.65%, KO: 18.45%) populations. The monocyte cluster was increased from 4.21% in WT tumors to 11.59% in *Klf4* KO tumors. DGE analysis (**Supplemental Figure VIA, Supplemental Table 14**), pathway analysis (**Supplemental Figure VIA&B, Supplemental Table 15&16**), and TF activity inference (**Supplemental Figure VIC**) showed macrophages and monocytes in *Klf4* KO samples exhibited reduced proliferation-related pathways and increases in inflammation related genes, pathways, and TF activities.

Thus, or data showed a major shift in the phenotype of tumor-associated macrophages (TAMs) in tumors from *Klf4* KO mice compared to WT mice. The macrophage phenotype in *Klf4* KO tumor-bearing mice resembled macrophages in naïve non-tumor-bearing lungs of WT mice supporting that altering the phenotype of AdvSca1-SM cells affects the phenotype of TAMs of the TME.

The compositions of Neutrophils (0.89% to 9.30%), B cells (1.19% to 3.20%), and T cells (3.95% to 8.69%) also exhibited increased percentages in AdvSca1-SM-specific *Klf4* KO tumors compared to WT tumors. We previously showed that LLC cells form immunologically “cold” tumors and are resistant to anti-PD1 therapy. To determine if there is a change in tumor-infiltrating T cells, IF staining for CD3^+^ T cells was performed to localize and quantify T cell infiltration in WT and *Klf4* KO tumors (**Figure 5**). Representative images (**Figure 5A**) and quantification (**Figure 5B**) indicate that compared to tumors in WT mice that exhibited low T cell accumulation, tumors from *Klf4* KO mice exhibited increased numbers of T cells within the core of the primary tumor as well as around the edges of the primary tumor. DGE analysis of T cells in the scRNA-seq data indicated elevated inflammation-related genes (**Supplemental Figure VIIA, Supplemental Table 17**), pathways (**Supplemental Figure VIIB&C, Supplemental Table 18&19**), and TFs activity (**Supplemental Figure VIID**), supporting immune activation of T cells in *Klf4* KO tumors. Collectively, our data support that AdvSca1-SM cells are a major component of lung tumors, significantly contributing through differentiation to CAFs. Furthermore, depletion of KLF4 in AdvSca1-SM cells, which changes the phenotype of AdvSca1-SM cells, inhibits tumor growth and alters the immune microenvironment, resulting in decreases in innate immunosuppressive macrophage populations, and increasing T cell infiltration into the tumor.

**Figure 5.**
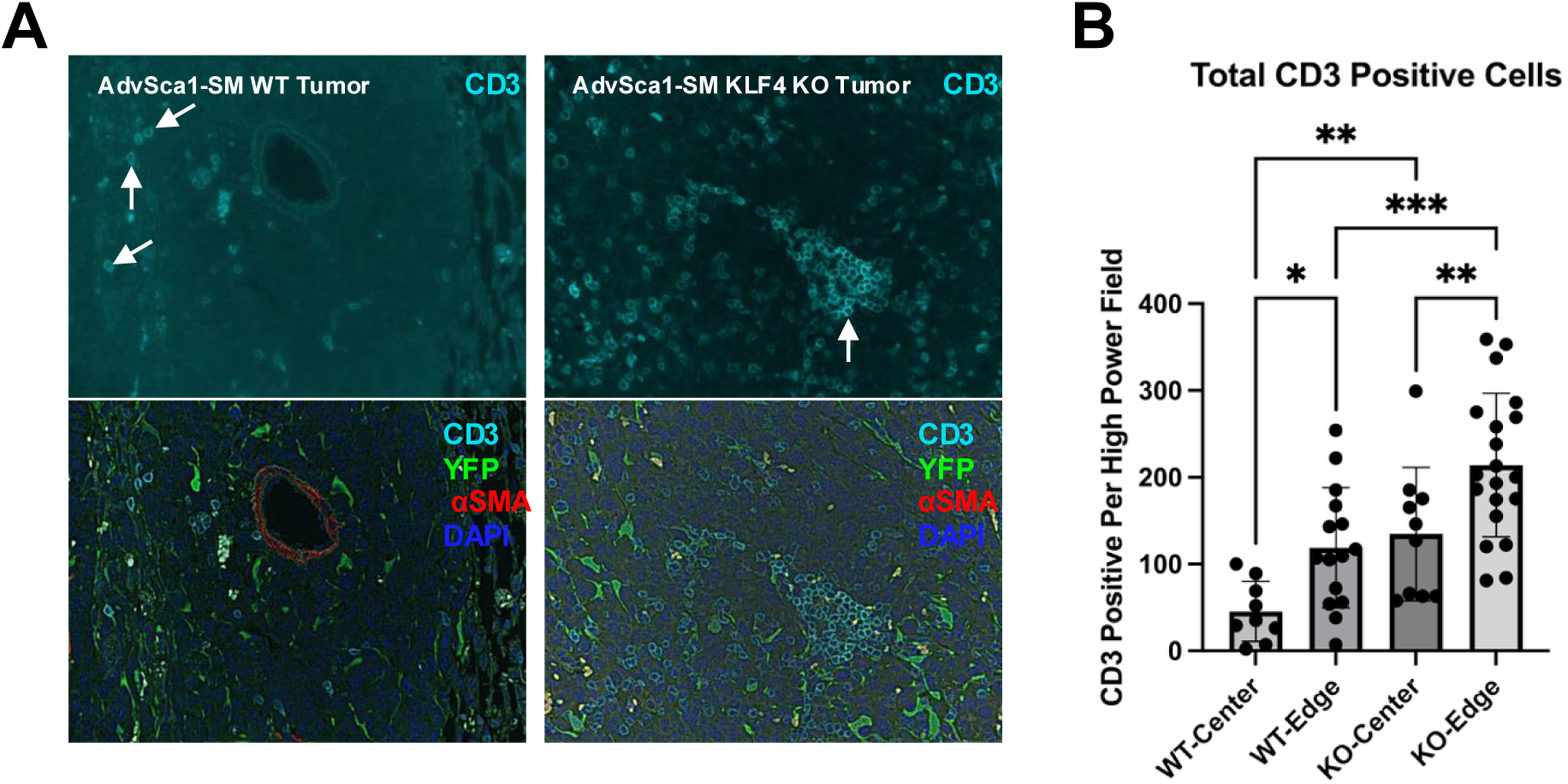
Increased T cell recruitment in primary tumors of AdvSca1-SM *Klf4* KO mice compared to WT mice. **(A)**. Lung sections from LLC tumor-bearing *Klf4* WT (**left panels**) and KO mice (**right panels**) were stained for CD3 (cyan), YFP (green; AdvSca1-SM lineage), aSMA (red), and DAPI (blue, nuclei) to identify recruited tumor-infiltrating T cells. Representative 40x images of CD3^+^ T cells in both the core and edge of tumors is shown **(B)** Cell count quantification revealed increased numbers of recruited T cells both in the core of the primary tumor as well as around the edges of the primary tumor. Arrows – representative CD3+ T cells. N=6; one-way ANOVA with Bonferroni’s post-hoc test; *p<0.05; **p<0.01; ***p<0.001.

To better decipher the cell-cell communication within tumors, we performed steady-state ligand-receptor inference and intercellular context factorization (ICF) using LIANA and Tensor- cell2cell. The analysis uncovers how cell communication changes between conditions and breaks down the complex cell-cell communization data into factors (factorization). We identified 9 factors, of which, Factor 5-9 exhibited significant changes in context loadings between naïve and tumor bearing lungs in WT mice (**Figure 6A**). Factor 5-7 exhibited higher context loadings in the tumor condition compared to naïve condition, suggesting that the gene expression pattern represented by these factors has a more prominent influence in the TME relative to naïve lung. Factors 8 and 9 exhibited reduced context loading in WT tumors compared to naïve lungs. The ICF analysis also quantifies the contribution of each cell type as a potential signal sender (**Figure 6B**) and receiver (**Figure 6C**). The results support that Factors 5 and 6 represent intercellular signaling from AdvSca1-SM cells, CAFs, and resident fibroblasts to endothelial cells (**Figure 6B&C, Supplemental Figure VIIIA**) and immune cells (macrophages, monocytes, neutrophils)(**Figure 6B&C, Supplemental Figure VIIID**), respectively. Further inspection of ligand-receptor pairs in Factor 5 indicated a strong signature of endothelial-to-ECM interaction (**Supplemental Figure VIIIB**). The reduction of Factor 5 in tumor-bearing lungs was attributed to decreased receptors involved in cell-cell and cell-matrix interactions by the endothelial cells (**Supplemental Figure VIIIC**). This data is consistent with tumor-induced changes in endothelial cell adhesion and migration. Factor 6 represents signaling from AdvSca1-SM cells to myeloid cells and their interaction with the ECM (**Supplemental Figure VIIIE**). The reduction in ligand-receptor interactions in factor 6 (**Supplemental Figure VIIIF**) indicated a potential shift of the myeloid cells towards a tumor-promoting phenotype (reduced migration (Cd44, Cd93, Plaur, Ccrl2) and phagocytosis activity (Cd36, Cd47). Factor 7 is characterized by the interaction between alveolar type I and II cells, endothelial cells, and immune cells (**Supplemental Figure VIIIH&I**). Overall, the decrease in Factor 5-7 likely indicated that tumors disrupted the normal cell-cell communization in resident lung cells, AdvSca1-SM cells, and recruited immune cells.

**Figure 6.**
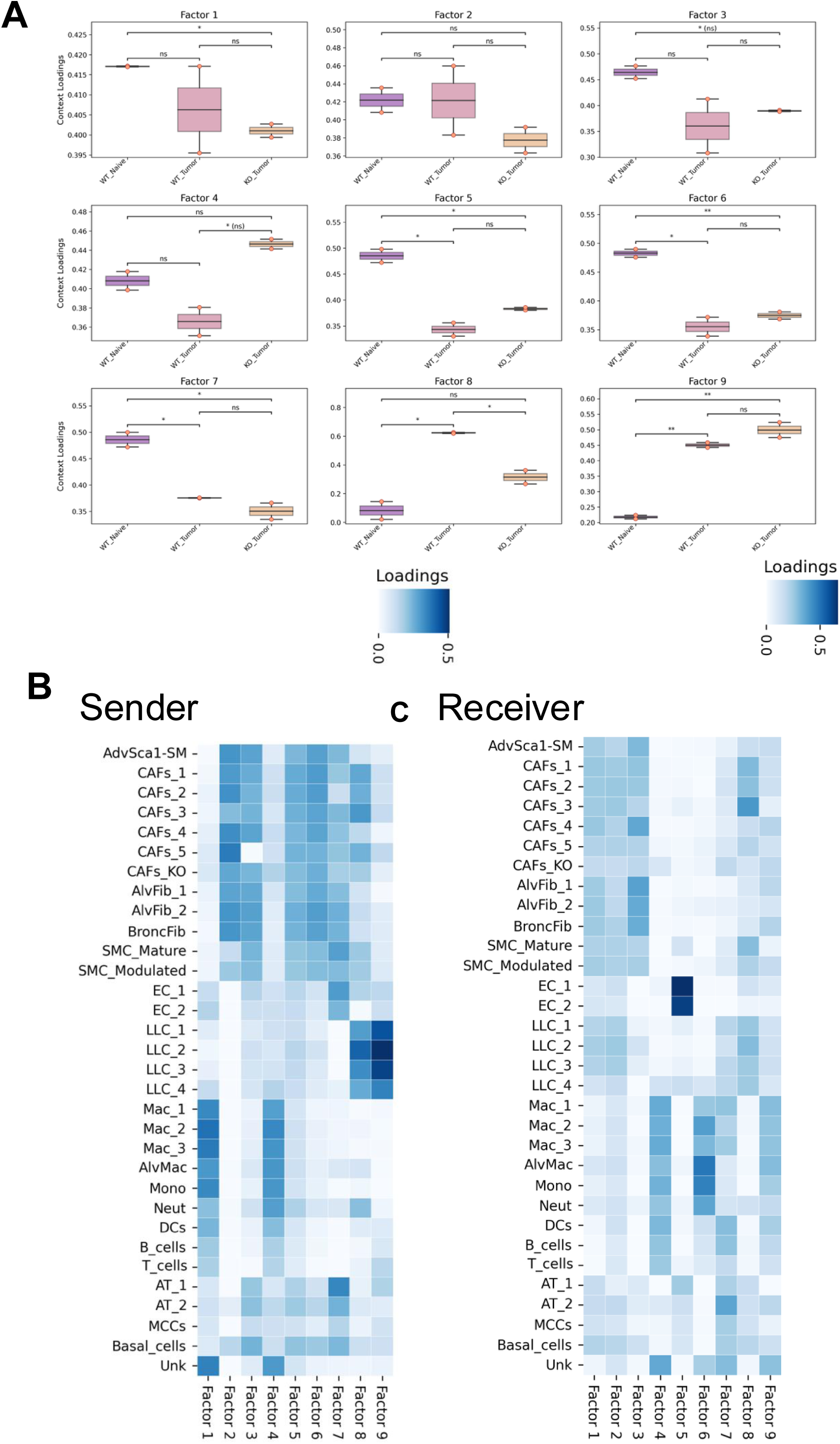
Cell-cell communication analysis reveals *Klf4* depletion in *Gli1*^+^ AdvSca1-SM lineage cells renders AdvSca1-SM-derived CAFs unresponsive to signaling from LLC cells. **(A)** Steady-state ligand-receptor inference and intercellular context factorization (ICF) was performed using LIANA and Tensor-cell2cell. Cell-cell communication was factorized into 9 factors, and their content loading were plotted between WT naïve, WT Tumor, and *Klf4* KO tumor conditions. Differences between groups was examined using independent t-test with false discovery rate correction (Benjamini-Hochberg method), as implemented by Tensor-cell2cell python package. ns not significant, *p<0.05, **p<0.01 Heatmaps depicting the sender clusters **(B)** and receiver clusters **(C)** loadings across intercellular communication factors is shown.

There was a significant increase in Factor 8 and Factor 9 interaction in tumor bearing lungs compared to naïve lungs (**Figure 6A**). Very interestingly, Factor 8 was suppressed in *Klf4* KO tumors compared to WT tumors (**Figure 6A**). Factor 8 represents communications from LLC tumor cells to CAFs (**Figure 6B&C, Supplemental Figure IXA&B**). Further examination of the ligand-receptor pairs (**Supplemental Figure IXB**) showed that Factor 8 represented ECM-to-cell interactions (collagens, proteoglycan, fibronectin to integrin receptors) and growth factor signaling (*Egfr, Fgfr1*). Select receptors indicated down-regulation in *Klf4* KO AdvSca1-SM cells and CAFs compared to WT (**Supplemental Figure IXC**), suggesting that *Klf4* depletion might induce the AdvSca1-SM-derived CAFs to a state that is unresponsive to the pro-tumorigenic signaling arising from LLC cells. Similarly, Factor 9 represented LLC-to-immune cell interactions (**Supplemental Figure IXD&E**) with the ligand-receptor pairs indicating immune cell recruitment and modulation.

## DISCUSSION

Alterations in the TME have are critical in regulating tumor progression and metastasis. The role of innate and adaptive immune cells has been extensively studied. Recently, the role of CAFs as a major functional component of the TME has been defined. CAFs represent a significant component of the TME in multiple cancers^35,36^. They produce a pro-tumorigenic ECM, a dynamic, non-cellular structure known to facilitate cancer progression and metastasis. CAFs communicate with cancer cells and are involved in crosstalk with other cells of the TME, including innate and adaptive immune cells. While accepted to be an important component of the TME and proposed to be central organizers of the TME, there are several questions that remain controversial, including their origin, the specific differentiation program(s) driving production of CAFs, the specific subtype(s) that predominantly contribute to tumor progression, and mechanisms mediating effects on cancer cells and other cells of the TME. Recent findings point to significant heterogeneity within CAF populations, yet the clinical impacts of distinct subtypes remain unclear. Several studies have indicated that specific CAF subtypes are associated with a poor prognosis. Distinct CAF populations have been identified in human LUAD with myofibroblastic CAFs (Myo CAFs)^37^ and HGF^Hi^/FGF7^Hi^ CAFs^38^ being correlated with poor overall survival.

However, there are major gaps in our understanding of the role of CAFs. Firstly, distinct subtypes of CAFs have been defined largely through the use of transcriptional programming.

What the role of each of these subpopulations is not clear. Secondly, the origin of CAFs remains unclear. It was long assumed that CAFs were derived from normal fibroblasts, in this case lung fibroblasts. From our data it is clear that a significant fraction of CAFs are derived from AdvSca1-SM cells that represent a stem/progenitor population derived from vascular smooth muscle. While AdvSca1-SM cells are present in the naïve lung in a stemlike phenotype, this population expands dramatically in the presence of tumor and undergoes differentiation into CAFs. From our data it is also clear that this population is a central regulator of tumor progression. Deletion of KL4F, specifically in this population, resulted in significantly smaller tumor growth. This was mediated through altering the phenotype of both the cancer cells as well as other cells of the TME and in particular immune cells. Analysis of cancer cell clusters showed enrichment of a slower proliferating subpopulation in tumors from *Klf4* KO mice (**Figure 7C**). In addition, subclustering the non-AdvSca1-SM-derived macrophage populations revealed a major shift in the phenotype of tumor-associated macrophages (TAMs) in *Klf4* KO mice compared to WT mice. The macrophage phenotype in KO tumor-bearing mice resembled macrophages in naïve non-tumor-bearing lungs of WT mice (**Figure 6F; black circles and Figure 7B**) suggesting that altering the phenotype of AdvSca1-SM cells affects the phenotype of TAMs of the TME. Finally, KLF4 deletion in AdvSca1-SM cells resulted in increased T cell infiltration into the tumor. Our scRNA-seq analysis did not identify large numbers of T cells.

This could be due the nature of the tumor; LLC tumors are resistant to immune checkpoint inhibitors, and are generally defined as “cold tumors”, with little T cell infiltration. Alternatively, our digestion procedure for the scRNA-seq analysis was not optimized for recovering this population, and future studies would attempt to enrich our samples for T cell populations.

While AdvSca1-SM cells can communicate with multiple cell types, the critical interactions controlling tumor growth remain to be defined. It is possible that factors produced by these CAFs directly promote cancer cell growth. Interestingly, CAFs derived from AdvSca1- SM cells produce HGF, and this is lost in *Klf4* KO CAFs. HGF produced by CAFs has been shown to be a paracrine factor for cancer cell growth^39^. However, in our hands, inhibiting HGF/MET signaling with crizotinib did not mimic the effects of *Klf4* knockout in CAFs. Further studies are needed to examine the state of MET activation in these models. Alternatively, CAFs may release factors that directly signal to macrophages, resulting in alteration of the macrophages to a more antitumor phenotype. Direct CAF interactions with T cells are also possible. To sort out these interactions will require depleting specific populations and determining if KLF4 deletion in the CAFs still reduces tumor growth.

The current studies do not define the trajectory of AdvSca1-SM cells during the initial stages of tumor development. Early scRNA-seq analysis using our orthotopic model may reveal a role for KLF4 in early tumor development. Alternatively, the lineage tracing mice could be crossed with genetic mouse models of lung cancer. In these models, intratracheal administration of Adeno-Cre knocks in lung-specific expression of oncogenic Kras, which subsequently results in tumor formation. These models recapitulate early tumor development and would provide insight as to the role of KLF4 in AdvSca1-SM cells during the transition from preneoplastic lesions to lung adenocarcinoma. Finally, all of our studies have been performed in Kras-driven lung cancer models. Studies during the past 15 years have subdivided lung adenocarcinoma according to oncogenic drivers. While the TME has been extensively studied in the context of immunotherapy in Kras driven lung cancer, less is known regarding components of the TME in lung cancers driven by mutations in receptor tyrosine kinases. Thus, investigating the role of AdvSca1-SM cells in these cancers remains to be examined.

In summary our studies indicate that AdvSca1-SM cells are a major source of CAFs in lung cancer, and that KLF4 which regulates the phenotype of AdvSca1-SM cells is an important regulatory control for cancer progression. Defining the downstream effectors of KLF4 may identify novel therapeutic targets to improve the response to standard of care therapies of lung cancer, and potentially other cancers.

## AUTHOR CONTRIBUTIONS

MCMWE and RAN designed the studies. SL, ACN, AMJ, TN, and AJJ performed the experiments. TN assisted with mouse colonies and microscopy imaging. SL performed the bioinformatics analysis. SL, RAN, and MCMWE wrote the manuscript. SL, ACN, AMJ, TN, AJJ, KAS, RAN, and MWM edited the manuscript.

## ACKNOWLEDGEMENTS

This work was supported by grants R01 HL 121877 (Weiser-Evans, Majesky), R01 HL151331 (Weiser-Evans) from the National Heart, Lung, and Blood Institute as well as R21 CA255246 (Weiser-Evans, Nemenoff) from the National Cancer Institute. Flow cytometry experiments were performed at the University of Colorado Cancer Center Flow Cytometry Shared Resource Core supported by Cancer Center Support Grant (no. P30CA046934). This resource is supported in part by the Cancer Center Support Grant (no. P30CA046934). The scRNA-Seq experiments were performed in conjunction with the University of Colorado Cancer Center Genomics and Microarray Shared Resource supported by Cancer Center Support Grant (no. P30CA046934).

## AUTHOR APPROVALS

All authors have seen and approved the manuscript, and that it hasn’t been accepted or published elsewhere. The manuscript has not been accepted or published anywhere else.

## LIST OF SUPPLEMENTAL TABLES

Supplemental Table 1: qPCR primers

Supplemental Table 2: DGE analysis comparing YFP^+^ cells in CAFs clusters versus AdvSca1- SM cluster (positive log2FoldChange indicate up-regulation in CAFs)

Supplemental Table 3: Pathway analysis of up-regulated genes in CAFs compared to AdvSca1- SM cells.

Supplemental Table 4: Pathway analysis of down-regulated genes in CAFs compared to AdvSca1-SM cells.

Supplemental Table 5: DGE analysis comparing Tumor associated macrophages and monocytes versus naïve (positive log2FoldChange indicate up-regulation in TAMs)

Supplemental Table 6: Pathway analysis of up-regulated genes in TAMs compared to naïve macrophages and monocytes.

Supplemental Table 7: Pathway analysis of down-regulated genes in TAMs compared to naïve macrophages and monocytes.

Supplemental Table 8: DGE analysis comparing YFP^+^ CAFs in *Klf4* KO versus WT samples (positive log2FoldChange indicate up-regulation in *Klf4* KO)

Supplemental Table 9: Pathway analysis of up-regulated genes in *Klf4* KO CAFs compared to WT.

Supplemental Table 10: Pathway analysis of down-regulated genes in *Klf4* KO CAFs compared to WT.

Supplemental Table 11: DGE analysis comparing LLCs in *Klf4* KO versus WT samples (positive log2FoldChange indicate up-regulation in *Klf4* KO)

Supplemental Table 12: Pathway analysis of up-regulated genes in *Klf4* KO LLCs compared to WT.

Supplemental Table 13: Pathway analysis of down-regulated genes in *Klf4* KO LLCs compared to WT.

Supplemental Table 14: DGE analysis comparing TAMs in *Klf4* KO versus WT samples (positive log2FoldChange indicate up-regulation in *Klf4* KO)

Supplemental Table 15: Pathway analysis of up-regulated genes in *Klf4* KO TAMs compared to WT.

Supplemental Table 16: Pathway analysis of down-regulated genes in *Klf4* KO TAMs compared to WT.

Supplemental Table 17: DGE analysis comparing T cells in *Klf4* KO versus WT samples (positive log2FoldChange indicate up-regulation in *Klf4* KO)

Supplemental Table 18: Pathway analysis of up-regulated genes in *Klf4* KO T cells compared to WT.

Supplemental Table 19: Pathway analysis of down-regulated genes in *Klf4* KO T cells compared to WT.

