## Supplemental Figures for "*Klf4*-dependent pivotal role of smooth muscle-derived *Gli1*^+^ lineage progenitor cells in lung cancer progression"

### Supplemental Figure I

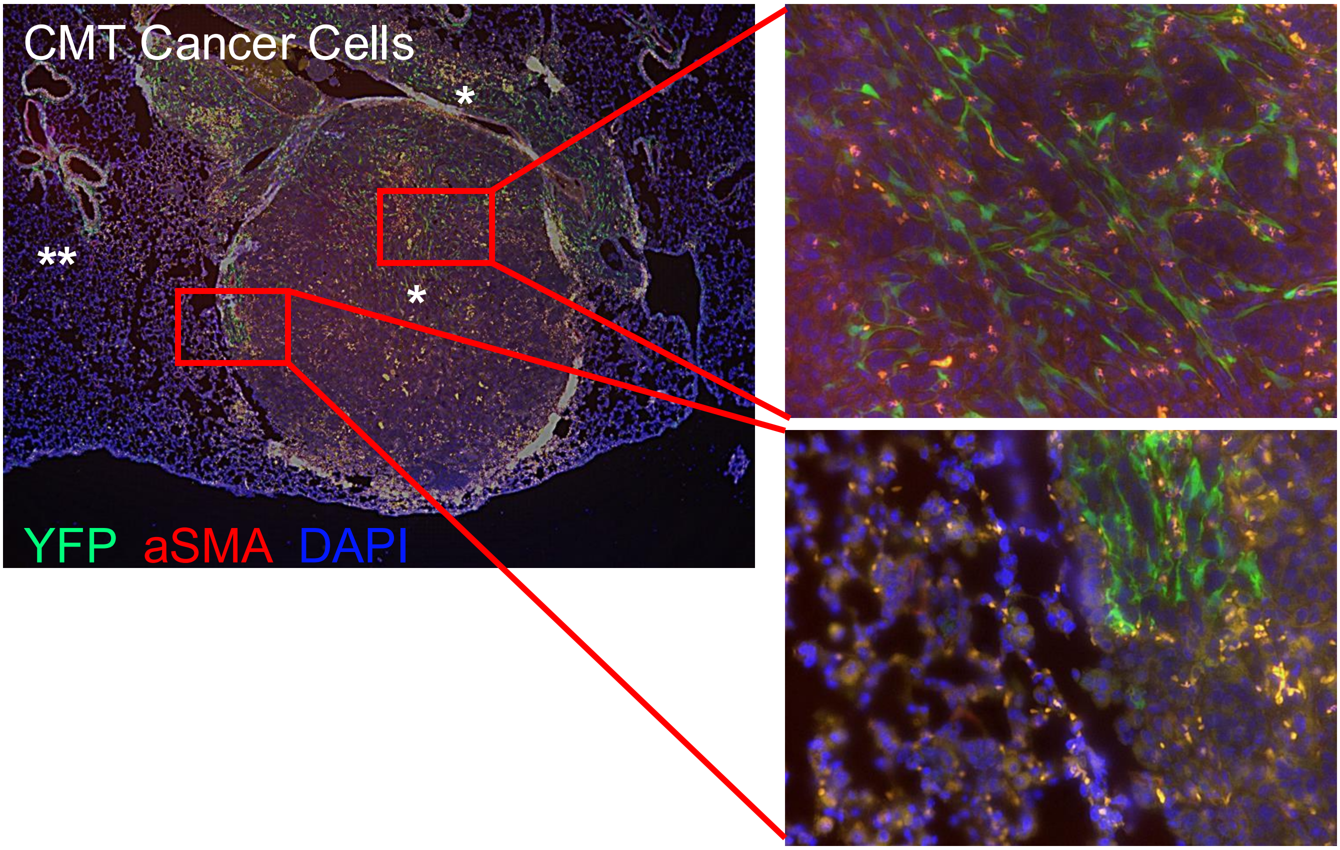


**Supplemental Figure I. *Gli1*+ AdvSca1-SM lineage cells expand and contribute to tumorigenesis in CMT KRas G12V tumors.** Tamoxifen-treated *Gli1*-Cre$^{ERT}$-YFP mice were injected with CMT167 cells orthotopically into the left lobe of the lung. Tumor-bearing lung tissues were immunofluorescently stained for AdvSca1-SM-derived cells (YFP, green), alpha smooth muscle actin (aSMA, red), and DAPI to identify cell nuclei. Right panels are higher magnification of boxed areas on the left showing AdvSca1-SM-derived cells in the core (**top right**) and periphery (**bottom right**) of the tumor. Representative 40x images shown on the right from N=4.

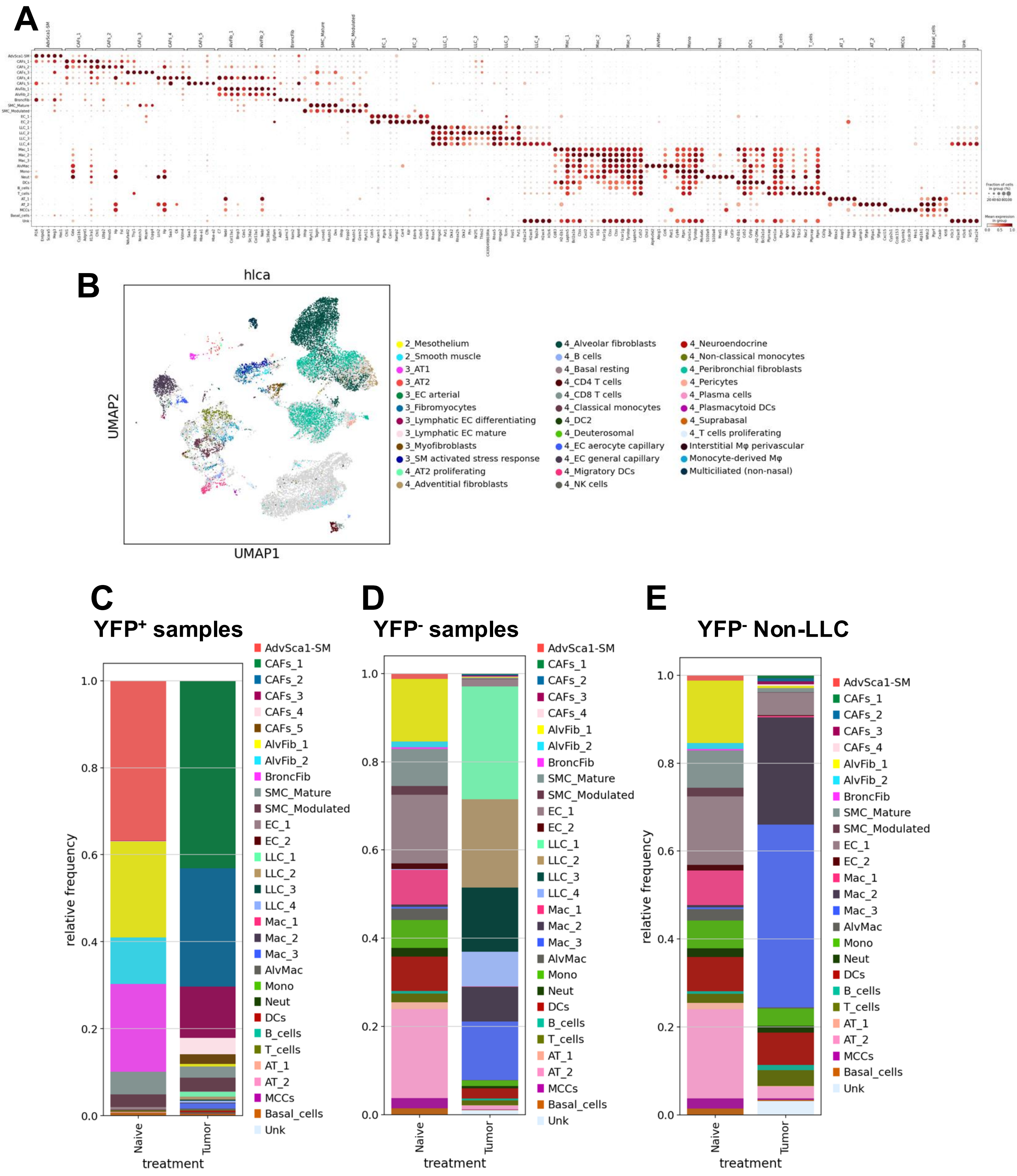


**Supplemental Figure II. scRNA-seq data annotation and composition analysis. (A)** Dotplot showing the top markers of each cluster from scRNA-seq data. **(B)** scRNA-seq data was mapped to the Human Lung Cell Atlas (HLCA) using homologous genes. Inferred cell identities based on HLCA alignment are show in the UMAP. Composition analysis was performed with YFP$^{+}$ **(C)**, YFP$^{-}$ **(D)**, and YFP$^{-}$ cells excluding tumor cells **(E)** and results shown in stacked barplots.

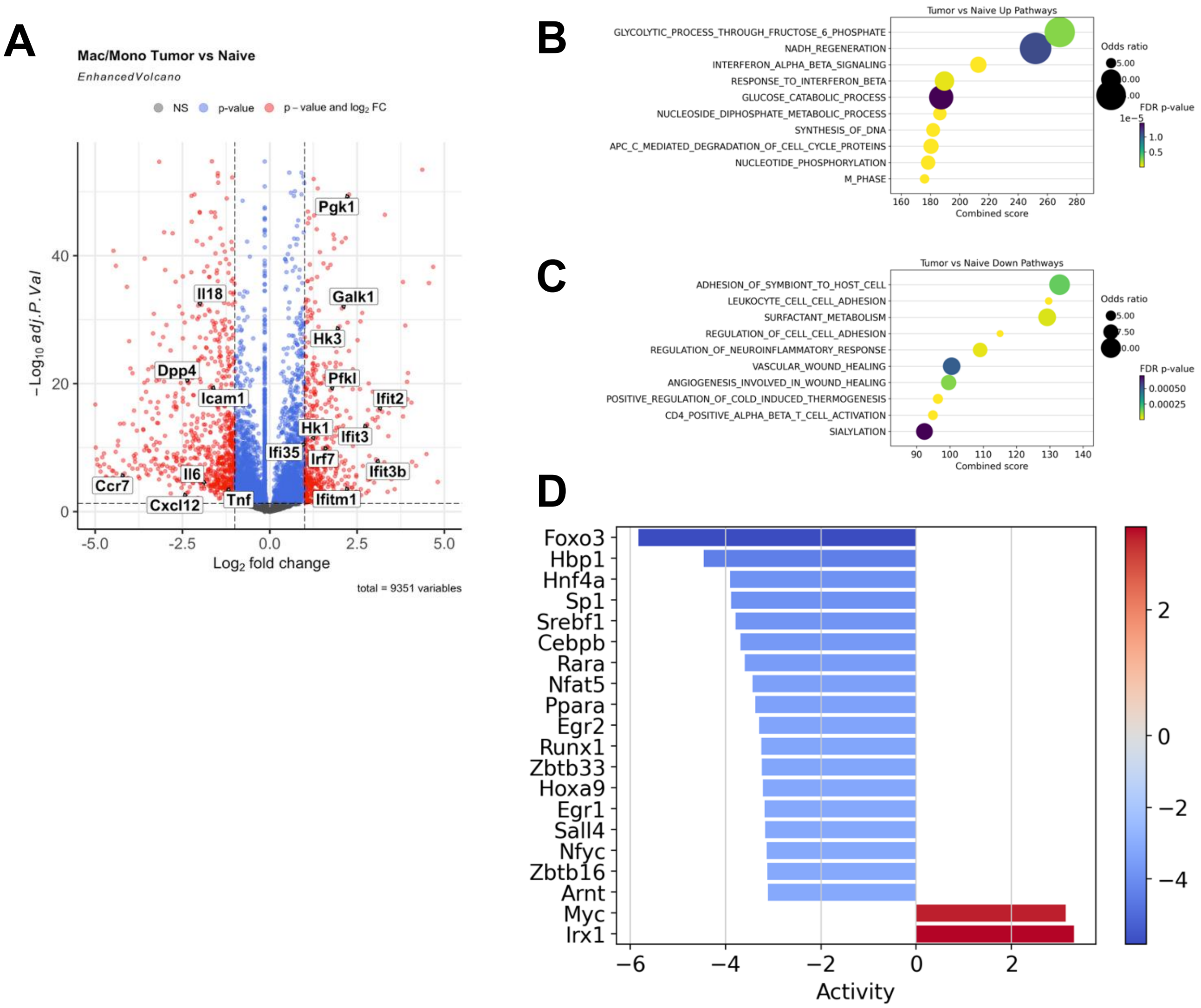


**Supplemental Figure III. DGE analysis between tumor-associated macrophages and monocytes and their naïve counterparts.** DGE analysis was performed with macrophages and monocytes in tumor-bearing and naïve lungs. **(A)** Volcano plot with select genes labeled. Positive logFC indicate up-regulation in tumor associated macrophages/monocytes. Pathway enrichment results with up-regulated genes **(B)** and down-regulated genes **(C)** in tumor-associated macrophages and monocytes are shown. **(D)** Transcription factors (TFs) activities were inferred based on differential gene expression analysis. Barplot shown with activated (red) and suppressed (blue) TFs.

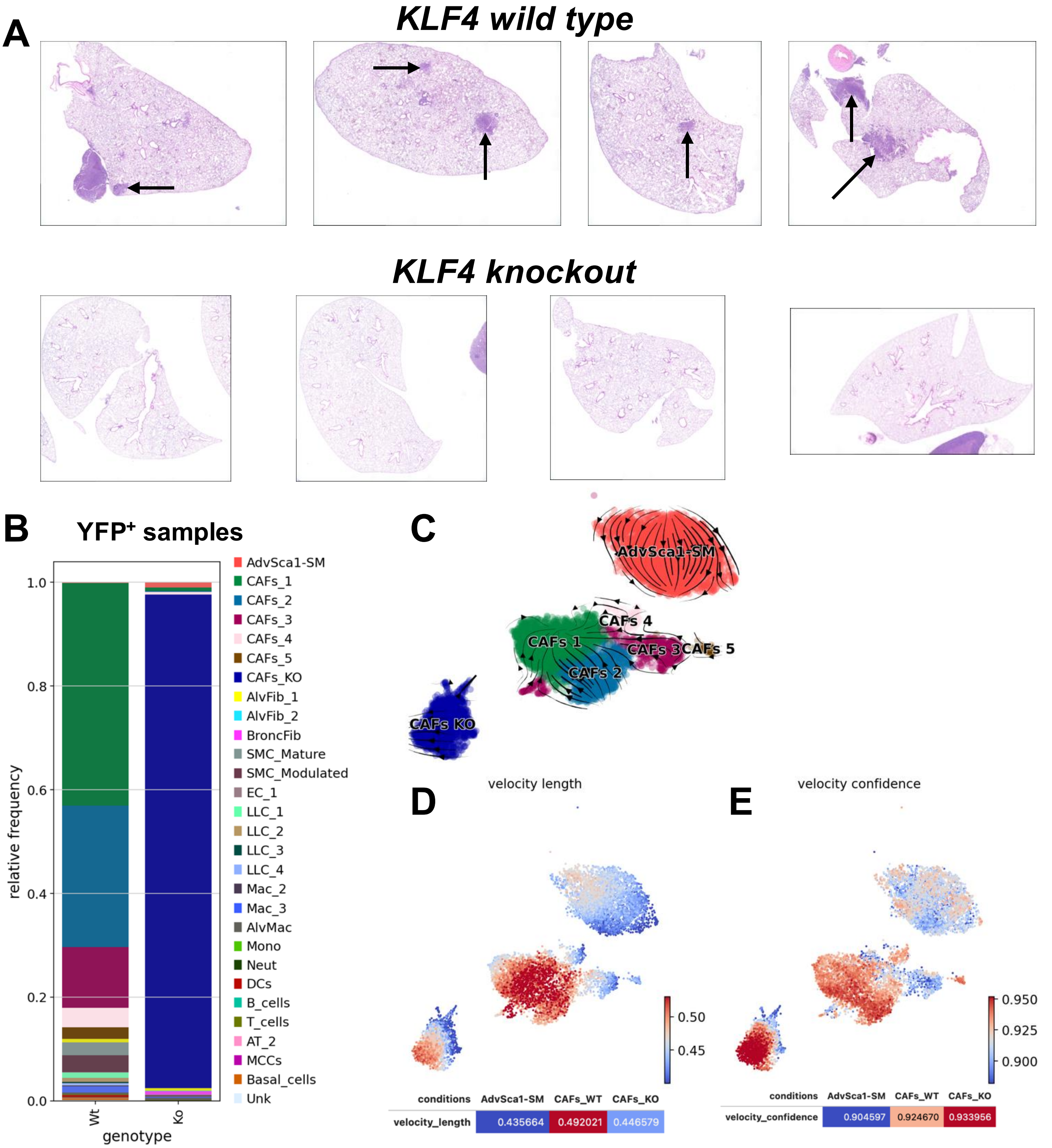


**Supplemental Figure IV. Composition analysis and RNA velocity analysis comparing *Gli1*⁺ AdvSca1-SM lineage cells in naïve and tumor bearing lung in WT and *Klf4* KO mice. (A)** Representative H&E images of tumor-bearing lung lobes from WT and *Klf4* KO mice showing secondary pulmonary metastases in WT, but not KO mice. Arrows=metastases. **(B)** Composition analysis was performed to contrast YFP⁺ cells from LLC tumor-bearing WT and *Klf4* KO lungs. Results are shown in stacked barplot. **(C)** RNA velocity analysis was performed between the naïve AdvSca1-SM cells, WT CAFs and *Klf4* KO CAFs clusters. Results are shown in stream plot. Velocity length **(D)** and velocity confidence **(E)** was plotted on UMAPs and quantified among AdvSca1-SM, WT CAFs and Klf4 KO CAFs clusters.

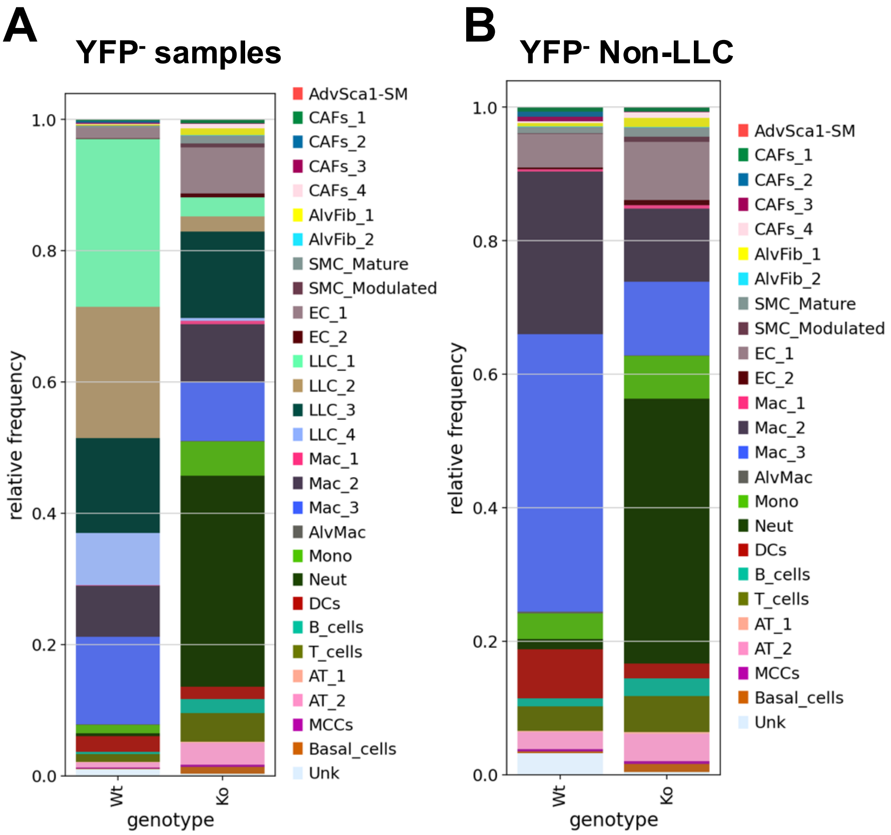


**Supplemental Figure V. Composition analysis comparing YFP$^{-}$ cells in tumor-bearing WT and *Klf4* KO lungs.** Composition analysis was performed with YFP$^{-}$ **(A)** and YFP$^{-}$ cells excluding tumor cells **(B)** and results are shown in stacked barplots.

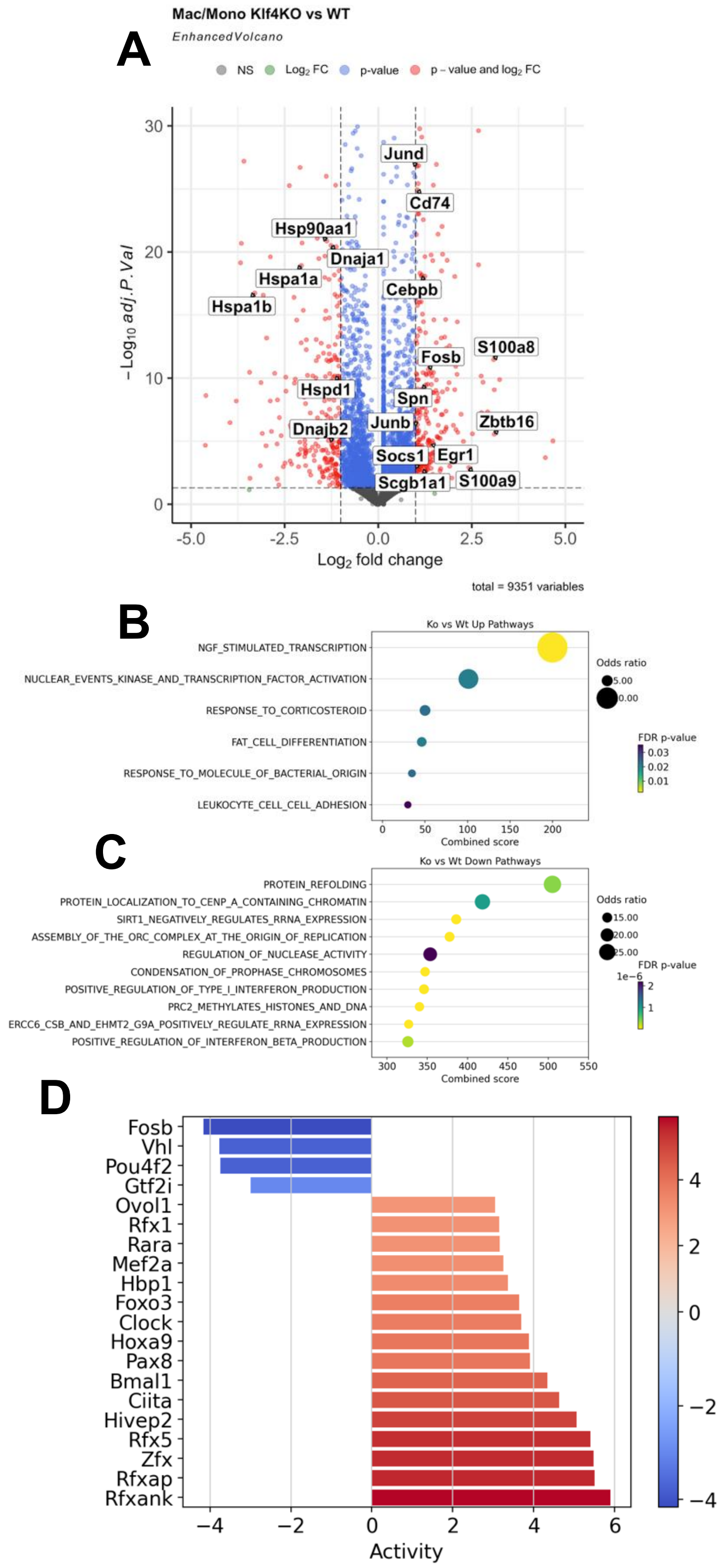


**Supplemental Figure VI. DGE analysis of tumor associated macrophages and monocytes comparing WT and *Klf4* KO samples.** DGE analysis was performed with macrophages and monocytes in tumor-bearing WT and *Klf4* KO samples. **(A)** Volcano plot is shown with select genes labeled. Positive logFC indicate up-regulation in *Klf4* KO. Pathway enrichment results with up-regulated genes **(B)** and down-regulated genes **(C)** in *Klf4* KO are shown. **(D)** TFs activities were inferred based on the differential gene expression analysis. Barplot is shown with activated (red) and suppressed (blue) TFs in *Klf4* KO samples.

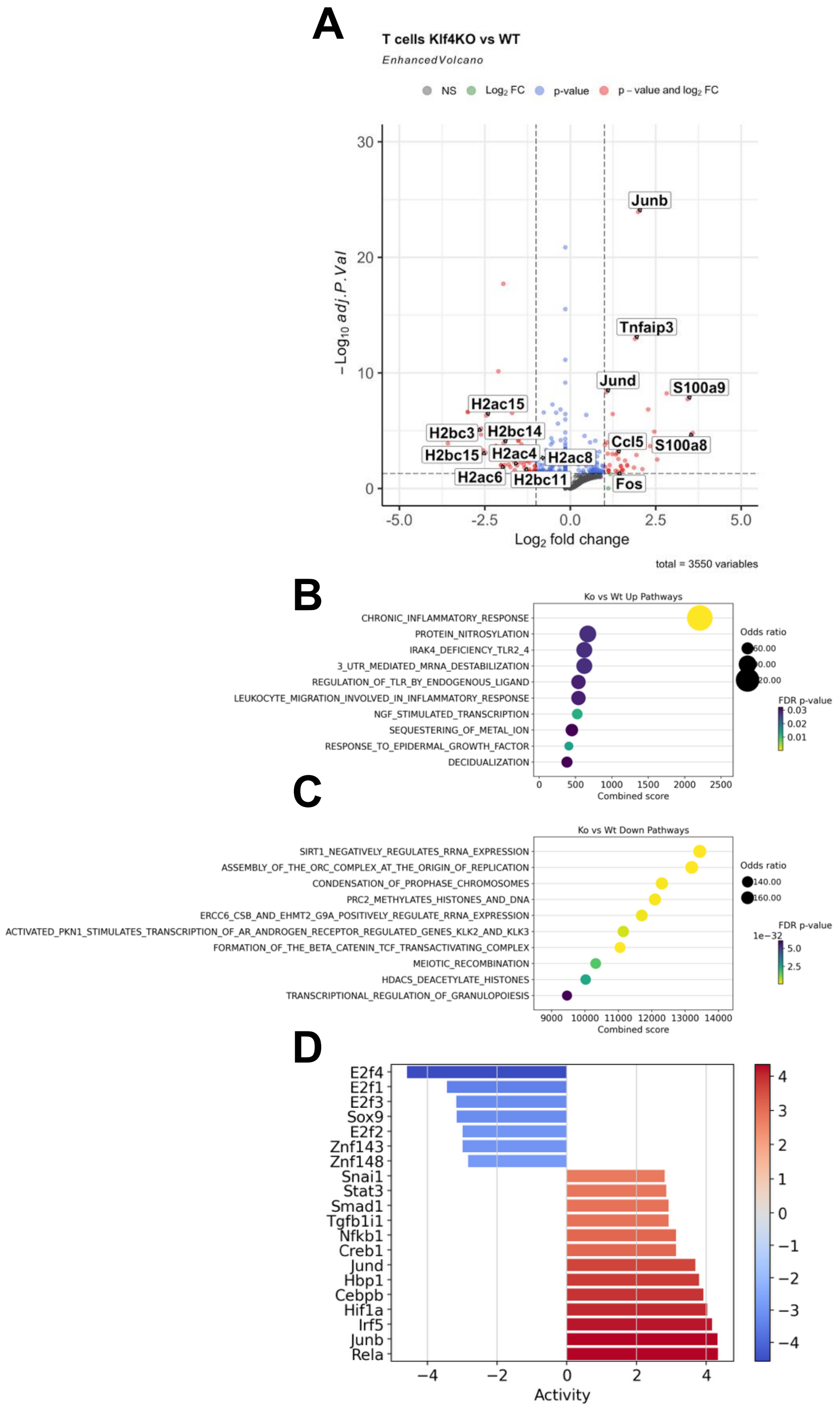


**Supplemental Figure VII. DGE analysis of T cells comparing WT and *Klf4* KO samples.** DGE analysis was performed with T cells in tumor-bearing WT and *Klf4* KO samples. **(A)** Volcano plot is shown with select genes labeled. Positive logFC indicate up-regulation in *Klf4* KO. Pathway enrichment results with up-regulated genes **(B)** and down-regulated genes **(C)** in *Klf4* KO are shown. **(D)** TFs activities were inferred based on the differential expression analysis. Barplot is shown with activated (red) and suppressed (blue) TFs.

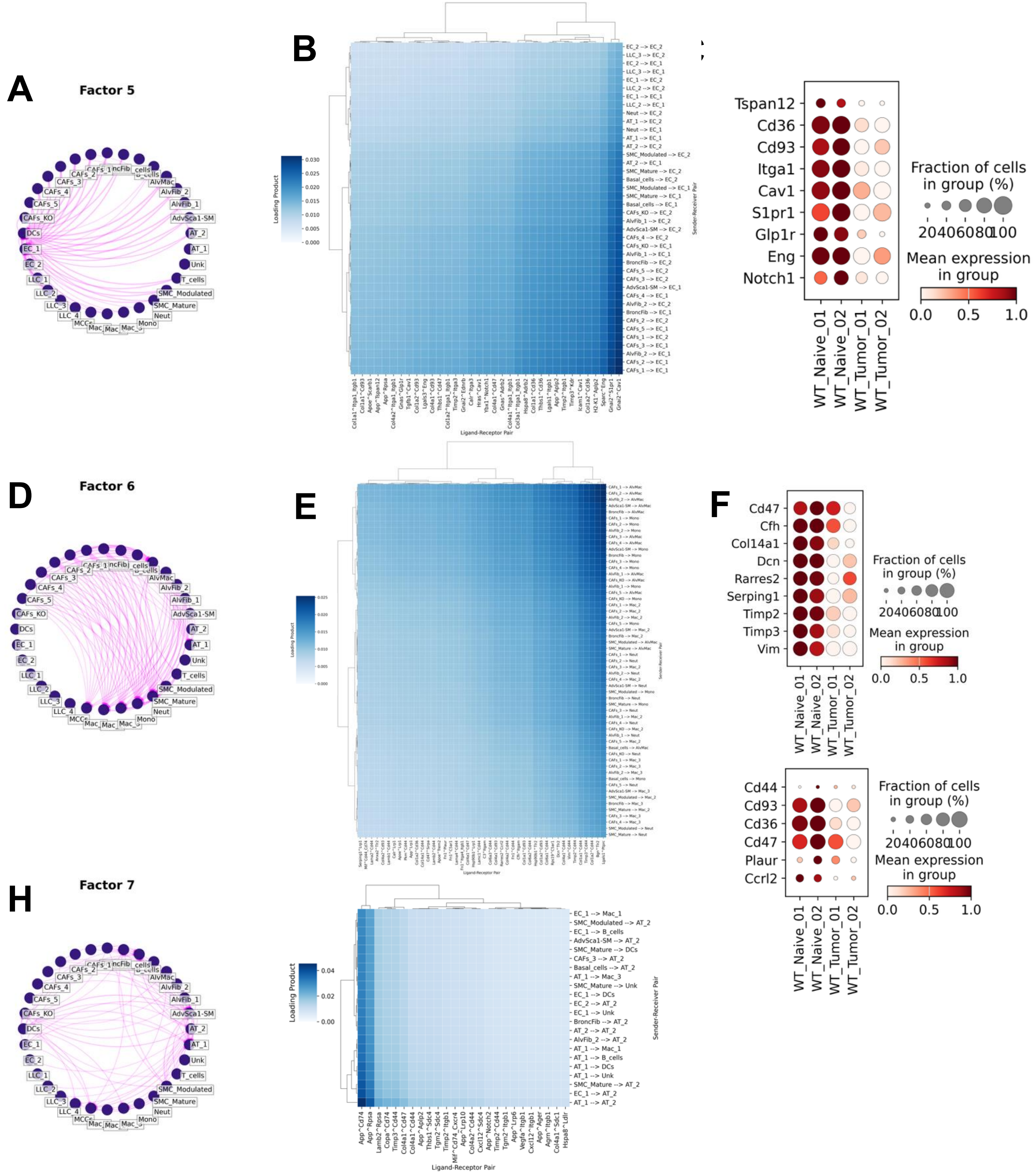


**Supplemental Figure VIII. Intercellular context factorization shows reduced cell-cell communication between CAFs, alveolar cells, endothelial cells, and immune cells in tumor-bearing lungs compared to naïve lungs.** Cell-Cell Communication (CCC) Network Plots **(A, D &H)** and heatmaps plotting sender-receiver pairs against ligand-receptor pairs **(B, E &I)** were generated for factor 5 **(A&B)**, 6 **(D&E)**, and 7 **(H&I)**. Dotplots were generated to show the expression of select receptors by endothelial cells in naïve and tumor-bearing lungs **(C)**, select ECM components by AdvSca1-SM cells, CAFs, and fibroblasts (**F**, top), and select receptors by immune cells (macrophages, monocytes and neutrophils)(**F**, bottom).

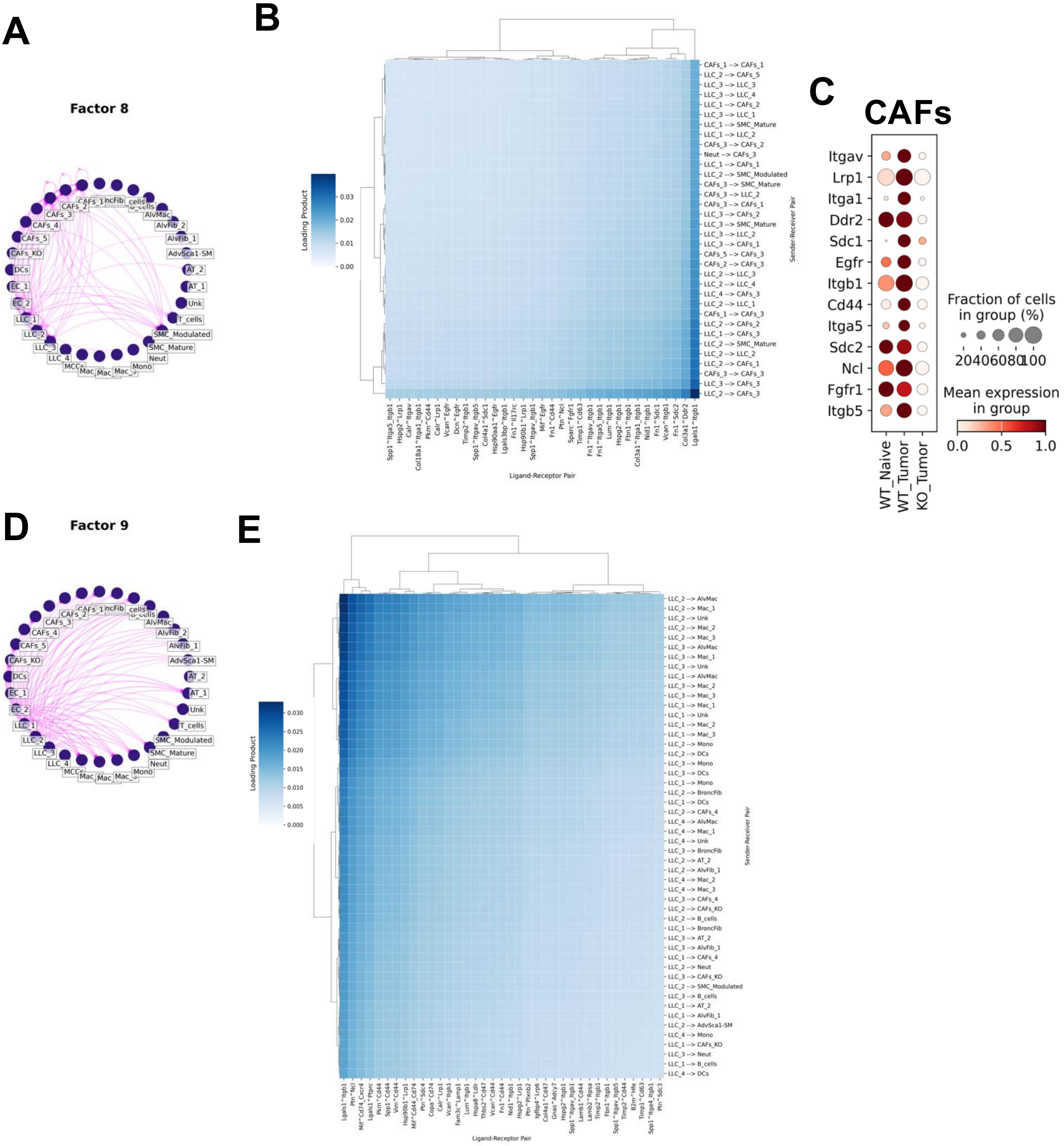


**Supplemental Figure IX. Intercellular context factorization showed reduced cell-cell communication between CAFs and LLC tumors in *Klf4* KO compared to WT lungs.** Cell-Cell Communication (CCC) Network Plot **(A&D)** and heatmap plotting sender-receiver pairs against ligand-receptor pairs **(B&E)** were generated for factors 8 **(A&B)** and 9 **(D&E)**. **(C)** Dotplot was generated to show the expression of select receptors by AdvSca1-SM cells and CAFs in *Klf4* KO and WT, naïve and tumor-bearing lungs.
